# Left/right asymmetry disruptions and mirror-image reversals to behavior and brain anatomy in *Ciona*

**DOI:** 10.1101/2021.03.03.433807

**Authors:** Matthew J. Kourakis, Michaela Bostwick, Amanda Zabriskie, William C. Smith

## Abstract

**Background:** Left-right asymmetries are a common feature of metazoan nervous systems. This is particularly pronounced in the comparatively simple larval central nervous system (CNS) of the tunicate *Ciona*, whose swimming tadpole larva shows a clear chordate ground plan. While common pathway elements for specifying the left-right axis are found in the chordates, particularly a requirement for Nodal signaling, *Ciona* differs from its vertebrate cousins by specifying its axis at the neurula stage, rather than at gastrula. Additionally, *Ciona*, and other ascidians, have a requirement for an intact chorionic membrane for proper left/right specification.

**Results:** We present here results showing that left-right asymmetry disruptions caused by removal of the chorion (dechorionation) are highly variable and present throughout the *Ciona* larval nervous system. While previous studies have documented disruptions to the conspicuously asymmetric sensory systems in the anterior brain vesicle, we document asymmetries in seemingly symmetric structures such as the posterior brain vesicle and motor ganglion. Moreover, defects caused by dechorionation include misplaced or absent neuron classes, loss of asymmetric gene expression, aberrant synaptic connectivity, and abnormal behaviors. In the motor ganglion, a brain structure that has been equated with the vertebrate hindbrain, we find that despite the apparent left/right symmetric distribution of interneurons and motor neurons, AMPA receptors are expressed exclusively on the left side, which equates with asymmetric swimming behaviors. We also find that within a population of dechorionated larvae, there is a small percentage with apparently normal left-right specification, and approximately equal population with inverted (mirror-image) asymmetry. We present a method based on a behavioral assay for isolating these larvae. When these two classes of larvae (normal and inverted) are assessed in a light dimming assay they display mirror-image behaviors, with normal larvae responding with counterclockwise swims, while inverted larvae respond with clockwise swims.

**Conclusions:** Our findings highlight the importance of left-right specification pathways not only for proper CNS anatomy, but also for correct synaptic connectivity and behavior.

## BACKGROUND

As sister taxa, tunicates and vertebrates each have a *central nervous system* (CNS) which shares important features: neurulation -- the folding of dorsal ectoderm into a tube, a tripartite brain, including a midbrain/hindbrain boundary and caudal nerve cord, and migrating neural crest or crest-like cells and neurogenic placodes, both of which contribute to sensory structures (Wada et al., 1998; Manni et al., 2004; Abitua et al., 2012; Stolfi et al., 2015; Abitua et al., 2015). Yet, the simple CNS of the swimming tadpole larva of the tunicate *Ciona* contains less than 200 neurons, a vast contrast to cell numbers in the millions or even billions for their nearest chordate relatives, the vertebrates (Nicol and Meinertzhagen, 1991). With an electron-micrograph derived synaptic connectome (Ryan et al., 2016), and a range of quantifiable behaviors (Bostwick et al., 2020a; Salas et al., 2018), the *Ciona* CNS is fertile ground for researchers seeking to bridge neural circuitry and behavior. Studies have begun to shed light on the circuit logic which governs larval behaviors. *Ciona* larvae have two distinct visuomotor behaviors, negative phototaxis and a looming shadow response, that are mediated by separate groups of photoreceptors, the Group I and Group II clusters, respectively (Kourakis et al., 2019). Both visuomotor circuits act via minimal circuits that consist of two-interneuron sequences. The photoreceptors first project to *relay neurons* (RNs in Figure 1) in the *posterior brain vesicle* (pBV). The relay neurons in turn project to the motor ganglion (MG), where they synapse to the *motor ganglion interneurons* (MGINs). The MGINs then synapse to the *motor neurons* (MNs). Intersecting with the looming shadow response is the negative geotaxis response. The antenna cells (Figure 1) sense the movement of the otolith pigment cell as the larva moves with respect to gravity, but the targets of the antenna cell in the pBV, the *antenna relay neurons*, are inhibited by photoreceptor RNs, unless the larva sees a light dim, at which point the inhibitions is released and the larva swims upwards (Bostwick et al., 2020a).

**Figure 1.**
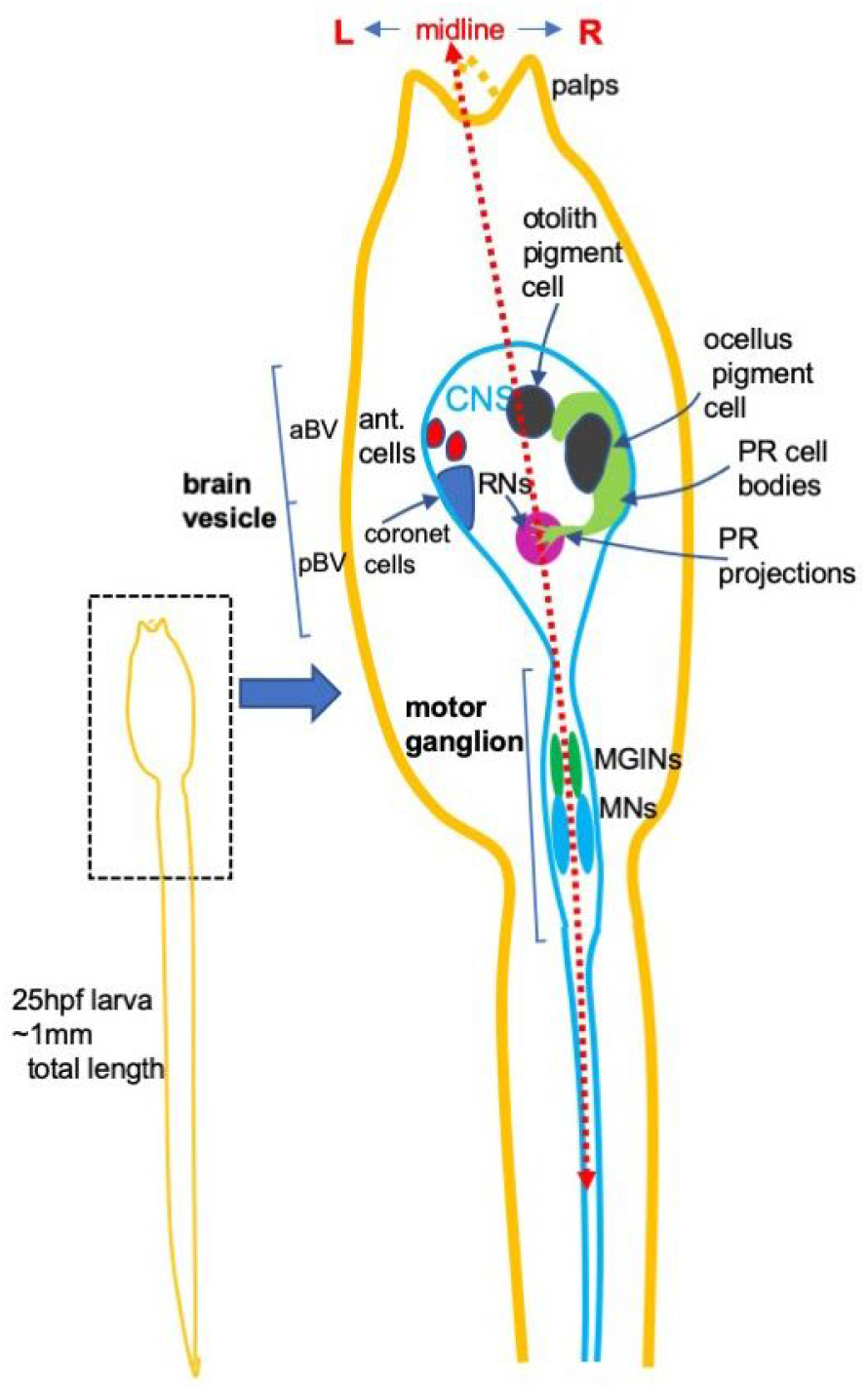
Cartoon of *Ciona* larva highlighting left/right asymmetry. Abbreviations: hpf: hours post fertilization; ant.: antenna; aBV: anterior brain vesicle; pBV: posterior brain vesicle; PR: photoreceptor; MGIN: motor ganglion interneuron; MN: motor neuron. L; left; R: right.

Most tunicates, including *Ciona*, are broadcast spawners - and hermaphrodites. The eggs are shed into the ocean enclosed in a chorionic membrane. The surface of the chorionic membrane is studded with maternal follicle cells, which may act to increase the buoyancy of the egg, while the chorionic membrane itself serves to prevent polyspermy and fertilization with self-sperm (Harada et al., 2008; Lambert, 2009). After approximately 20 hours of development (Matsunobu and Sasakura, 2015) the newly-formed *Ciona* larva will hatch from the chorionic membrane and start swimming. Most experimental procedures for the *Ciona* egg and embryo require the chemical removal of the chorion. These procedures include *in situ* hybridization, most microinjection protocols, CRISPR gene editing, and the widely used electroporation method for transgenesis. Moreover, *in situ* hybridization in larvae also requires the earlier removal of the chorion. This is due to the presence of a group of maternal cells, known as the test cells, that occupy the space between the egg surface and the chorionic membrane. As the embryo develops, the test cells adhere to the epidermis where they contribute to the test, a cellulose-containing sheath that encapsulates the larva (Satoh, 1994, 2014). The larval test impedes the efficient penetration of *in situ* hybridization probes and antibodies, and itself can give very high background staining. Fortunately, dechorionation removes both the chorionic membrane and the test cells, allowing for *in situ* hybridization of larvae. Most tunicate researchers perform dechorionation, either with trypsin or Na-thioglycolate/protease-based solutions, before fertilization or, alternatively, immediately after, but before the first cleavage plane has formed (Christiaen et al., 2009; Hendrickson et al., 2004; Veeman et al., 2011).

Despite the usefulness of this procedure, there are well-documented disruptions caused by early dechorionation to the left-right asymmetry of tunicate embryos and larvae (Boorman and Shimeld, 2002; Shimeld and Levin, 2006). A Nodal-dependent left-right asymmetry has been observed broadly across the chordates (and deuterostomes more generally), and generates the internal asymmetry of, for example, endodermal organs and heart in vertebrates (Duboc and Lepage, 2008; Soukup et al., 2015; Hamada and Tam, 2020). *Ciona* likewise shows internal Nodal-dependent asymmetries (Tanaka et al., 2019; Yoshida and Saiga, 2008, 2011). However, unlike vertebrates where asymmetric expression of downstream targets such as Pitx and Lefty are established during the gastrula stage, tunicates show a critical period for establishment of asymmetry at the neurula to early tailbud stages (Shimeld and Levin, 2006). Importantly, the presence of the chorionic membrane is required in tunicates up to the early tailbud stage for proper establishment of left-right asymmetry. Drugs such as proton pump inhibitors have similar effects as early dechorionation, as neurula-stage embryos treated with these drugs show altered symmetry of internal organs after metamorphosis (Shimeld and Levin, 2006; Kourakis and Smith, 2007) suggesting that the removal of the chorionic membrane may work, in part, through disruption of the ionic environment (Shimeld and Levin, 2006). Contact between the left epidermis with the inner chorionic membrane appears to be required for left-sided *Nodal* expression in *Halocynthia* (although this may vary across ascidian species) (Nishide et al., 2012; Yamada et al., 2019). Consistent with this requirement, the application of pressure so that both left and right epidermis contact the chorion results in bilateral *Nodal* expression (Nishide et al., 2012).

Among the structures with left-right asymmetry in *Ciona*, the larval CNS is particularly pronounced (Boorman and Shimeld, 2002; Oonuma et al., 2016; Ryan et al., 2016). Figure 1 summarizes the gross asymmetries in the *Ciona* larval CNS. These include the larval visual organ, the ocellus, which is located on the right side of the brain vesicle, and the otolith-associated antenna cells on the left side, which, along with the otolith pigment cell comprise the gravity-sensitive otolith organ. Also asymmetrically located in the ventral portion of the left brain vesicle are the 15 coronet cells, which have been speculated to be components of a sensory system, but of unknown function. The coronet cells are dopaminergic (Moret et al., 2005), and receive their name from the coronet cells of the saccus vasculosus, an organ that serves as a photoperiodic and seasonal sensor in teleosts (Nakane et al., 2013). Many other examples of left-right asymmetry are evident from the *Ciona* synaptic connectome, including left-right asymmetric projection of axons (Ryan et al., 2016).

We show here that the disruptions to left-right asymmetry caused by dechorionation extend beyond the sensory systems and are observed throughout the length of the larval CNS. Moreover, the disruptions vary extensively from larva to larva. Disruptions are observed not only to the cellular anatomy of larvae, but to the connectivity of neurons and to larval behavior, as well. While the anatomy and behavior of early-dechorionated larvae on average was highly abnormal, we find that among dechorionated larvae a subpopulation with apparently more normal CNS development and behavior can be identified, as well as a subpopulation of larvae with mirror-image CNS asymmetry and behavior.

## MATERIALS AND METHODS

### Animal care

*Ciona robusta* adults were collected from Santa Barbara Harbor and kept at the University of California, Santa Barbara marine laboratory. Single clutches of larvae were obtained by collecting and mixing eggs and sperm from three adults, placing the zygotes at 18°C until they had reached the desired stage (Veeman et al., 2011).

### Dechorionation

Embryos were dechorionated using the sodium thioglycolate/protease method, as described previously (Christiaen et al., 2009). For early dechorionation, eggs are incubated at room temperature in the dechorionation solution, beginning 2 minutes after fertilization. Embryos are washed in seawater 5x in 60 mm 1% agarose-coated plastic petri dishes, then moved to 18°C until the desired time post-fertilization is reached. For late dechorionation, the dechorionation solution is added to embryos at the late-tailbud stage, about 14 hours post fertilization (hpf) (Hotta et al., 2007). Embryos are then washed in seawater, as above, and placed at 18°C in agarose-coated dishes. Following dechorionation, whether early or late, left-right sidedness of asymmetric features was always determined with reference to the dorsal midline.

### Immunohistochemistry and *in situ* hybridization

Antibody labeling was carried out as follows: *C. robusta* larvae were fixed in sea water + 2% paraformaldehyde for 1 hour at room temperature, before washing three times in PBS + 0.1% Triton X-100 (PBT). Larvae were placed in PBT + 5% lamb serum for 1 hour at room temperature and transferred to PBT + 5% lamb serum with a *C. robusta* Arrestin antibody raised in rabbit (gift of Takehiro Kusakabe), at a dilution of 1:1000, overnight at 4°C. Following five washes in PBT at room temperature, a secondary antibody, α-rabbit AlexaFluor 488 (Invitrogen), was used at 1:1000 overnight at 4°C, followed by five more PBT washes at room temperature before imaging.

*In situ* hybridization was carried out as previously described using the hybridization chain reaction (hcr) v. 3.0 protocol of Molecular Instruments (https://files.molecularinstruments.com/MI-Protocol-HCRv3-GenericSolution-Rev6.pdf). RNA probe sets were designed to the following *Ciona* genes: tyrosine hydroxylase (TH), vesicular glutamate transporter (VGLUT), vesicular GABA transporter (VGAT), Pitx, AMPAR, and vesicular acetylcholine transporter (VACHT). Larvae were imaged on Olympus Fluoview1000 or Leica SP8 confocal microscopes. Post-acquisition analysis and rendering used Imaris v.9 or ImarisViewer v.9.5.1 software.

### Behavioral assays

Dechorionated larvae were raised in 0.2% methyl cellulose in filtered sea water to prevent them from sticking to each other. Any unfertilized eggs, or larvae with gross morphological defects *(e*.*g*., kinked tails, open brains), which can occur as a result of damage to eggs during dechorionation, were removed before behavioral assays. Gravitaxis and dimming-response behavioral assays were performed as described previously (Bostwick et al., 2020a; Kourakis et al., 2019), except dishes were always coated with 1% agarose and larvae were assayed in 0.2% methyl cellulose in filtered sea water. All larval behavioral assays were performed at 25 hours post fertilization. For these assays, we waited 5 minutes between movies to allow larvae to recover. During all gravitaxis assays, the 505 nm LED was dimmed from 600 to 0 lux and it was positioned above the petri dish [top illumination condition, (Bostwick et al., 2020a)], shining down onto the dish at a 45 degree angle. Movies for gravitaxis assays were recorded for 20 seconds prior to the dim and 20 seconds post-dim.

To assess the left/right asymmetry of dimming-response swim trajectories (*i*.*e*., clockwise or counterclockwise), larvae in 6 cm petri dishes with 0.2% methylcellulose/seawater were recorded from above with 700 nm illumination and exposed to a 505 nm LED dimmed from 600 or 1500 to 0 lux.

### Selection of photo- and gravity-responsive dechorionated larvae

Dechorionated larvae at 23-24 hpf were placed in a 1% agarose-coated 6 cm petri dish and observed under a dissection microscope. Room lights were turned off and a fiber optic light source with two stems was used to illuminate the sample dish from left and right sides. A field of view was chosen containing at least several larvae, taking care to remove any larvae at the air-water interface; if larvae were found at the interface, they were removed with a BSA-coated pipet tip and placed at the bottom of the dish. At intervals of approximately every half minute, the fiber optic light source was dimmed, either by switching off the light at the source or by covering the fiber optic stems with the observer’s thumbs or fingers. After ∼5 seconds, light was reintroduced and the field of view observed. If any larvae were seen swimming upwards towards the surface in response to the dimmed light, they were removed and placed in a separate agarose-coated petri dish for later behavioral assay or for fixation and *in situ* hybridization (see above). The larvae selected in the upward swimming assay were compared to larvae from the same dish that did not show upward swimming behavior, but were judged to have good overall head-to-tail morphology and a tail-flick response to being touched, as an indication of a functioning motor system. Sibling larvae that were not dechorionated were also kept as controls. The same procedure was followed to find larvae for the swim reversal assay, but after showing upward swimming following dimming, the larvae were sorted by their pigmentation asymmetry, separating those with right-sided ocelli from those with left-sided ocelli. Larvae were placed in a 6 cm 1% agarose-coated petri dish containing seawater with MS-222 at 50 mg/mL, a concentration at which larvae swam more slowly, allowing for manual sorting by pigmentation asymmetry. Larvae were in the sorting solution at most 5 minutes, after which time they were transferred to a separate dish with seawater.

### Quantification of behavior/statistical analysis

Behaviors were assayed from at least two different clutches of larvae for each experimental condition, and two or more movies were collected from each clutch. However, we were only able to collect one movie from one of the three non-dechorionated larvae clutches due to an accident in which the petri dish broke after the first movie. Movie editing and behavior analyses were aided with the ImageJ software package. Gravitaxis behavior was evaluated by selecting ∼20 larvae at random from each recording and manually tracking their post-dim swims over 20 seconds, based on previously described criteria (Bostwick et al., 2020b). A dim response was qualified as any initiation of swimming immediately following dimming. From each group of recordings for a given condition, the percentages of larvae responding to the dim and swimming in each direction were calculated. Swims were assessed in the 20 seconds following dimming, and categorized as UP, sideways (SIDE) or DOWN, and the percentage in each category plotted in 45 degree wedges (Figure 7A and B). Swim directions were assigned a value (down = 1, sideways = 2, up = 3). Swim speed and tortuosity were calculated using a MATLAB script (Kourakis et al., 2019). Tail beat frequencies (flicks per second) were manually scored using 20 second clips. Comparison of behavioral results between conditions was done as described previously (Bostwick et al., 2020b), using a Wilcoxon Sum Rank Test (Mann Whitney U Test) to assess statistical significance.

Time-lapse recordings to assess left/right asymmetry of swimming were evaluated by manually tracking the larvae over 13 seconds following dimming of the 505 LED lamp, and then scoring the trajectory of each swim as clockwise, counterclockwise or straight. Each larvae was assessed three to five times, with 5 minute recovery periods between dimming trials.

## RESULTS

### Profound and highly variable defects to the pigmented sensory systems caused by early dechorionation

Early dechorionation of tunicates is known to give developmental defects, including to the CNS (Yoshida and Saiga, 2011). However, the extant and range of developmental abnormalities resulting from early dechorionation has not been fully characterized. The pigmented cells of the brain vesicle ocellus and the otolith sensory organs are perhaps the most easily assayed asymmetries in the *Ciona* body, imaged here with a compound microscope, but visible even under a simple dissecting scope (Figure 2). The pigments cells are critical for the proper functioning of these sensory organs (Jiang et al., 2005; Tsuda et al., 2003). The otolith contains a single large melanized pigmented cell attached to the ventral surface of the brain vesicle. The melanin appears to contribute to the function of the pigment cell by increasing its specific gravity by chelating ions such as potassium, calcium and zinc (Sakurai et al., 2004). When a larva changes its orientation with respect to gravity, the movement of the pigment cell within the ventricle is sensed by the antenna cells. The pigmented cell of the ocellus directionally shades the Group I photoreceptor outer segments, and allows a larva to determine the direction of light by performing casting swims (Salas et al., 2018). In the non-dechorionated *Ciona* larva the otolith pigment cell appears nearly spherical and is found near the midline (Figure 2A). By contrast, the pigment cell of the ocellus forms a shallow cup and is found on the right side of the brain vesicle, slightly posterior to the otolith.

**Figure 2.**
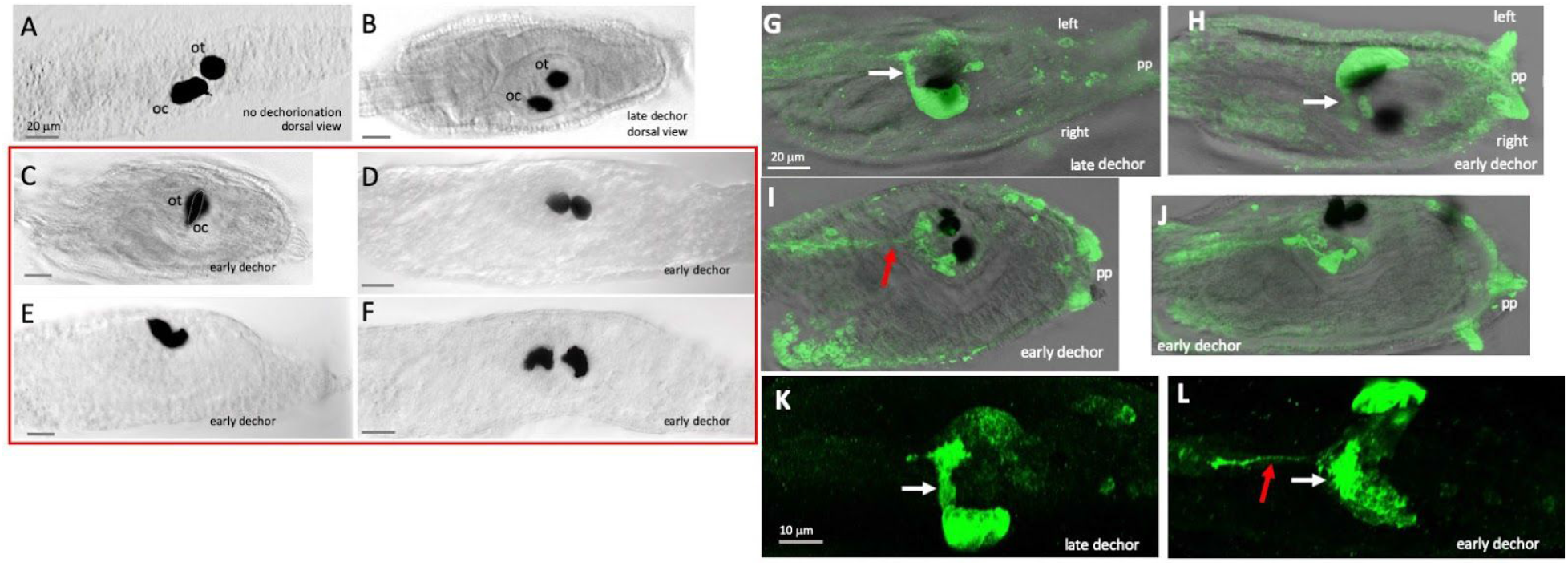
Disruptions to the ocellus caused by early and late dechorionation. **A** and **B**. Pigment cells in representative non-dechorionated and late-dechorionated larvae, respectively. **C**-**F**. Representative larvae showing a range of pigment cell defects in early-dechorionated larvae, including: overlapping otolith and ocellus at the midline (ocellus is outlined) **(C)**; two pigmented structures both resembling otoliths **(D, see also I)**; a single fused pigment spot **(E)**; and two pigmented structures, both resembling ocelli **(F). G-L**. Photoreceptors in early and late-dechorionated larvae immunostained with anti-Arrestin. G. late-dechorionated larva. **H**. Early-dechorionated larva showing left/right reversal of photoreceptors. **I-J**. Examples of early-dechorionated larvae with abnormal numbers and locations of photoreceptors. **K** and **L**. Late and early-dechorionated larvae, respectively, showing apparently normal photoreceptor development in late-dechorionated larva, while the in early-dechorionated larva the photoreceptor cluster is reversed, and while most projections are targeted correctly, one or more is projecting posteriorly. White arrows indicate properly targeted photoreceptor projections, and red arrows indicate aberrant photoreceptor projections. Abbreviations: ot: otolight; oc: ocellus; pp: palp; decor: dechorionation.

To determine what effect dechorionation had on the otolith and ocellus pigment cells, we dechorionated embryos either early or late (one cell and late tailbud stages, respectively) and then assessed 25 hpf larvae for pigment cell morphology. Control larvae whose chorions were not removed showed the expected phenotype for pigmented cells (Figure 2A) (38 normal of 40 total; of the two larvae remaining, one showed otolith and ocellus pigment cells at the midline, while another showed split/double otolith pigment clusters with an otherwise normally positioned ocellus). Late-dechorionated larvae showed the expected pigmented cell phenotype and asymmetry in all cases (28 normal/28 total). By contrast, when embryos were dechorionated early, a range of abnormalities were observed in the resulting larvae with respect to the non-dechorionated and late-dechorionated larvae. Of 34 early-dechorionated larvae examined, only 9 had cells that would be classified as an otolith and an ocellus pigment cell based on their morphology *and* that showed proper left-right asymmetry; of the remainder, eight had mirror-image reversals of the pigment cells. In fifteen larvae, we observed both pigmented cells positioned at the midline (*e*.*g*., Figure 2C, I). In the remaining nine larvae the pigmented cells either lacked the characteristic spherical or cup-shaped morphologies, or displayed an unexpected combination of morphologies, suggesting that in some cases two otolith or ocellus pigment cells had formed (see Figures 2 C-F for a range of pigment cell morphologies). We also observed a single larva which lacked pigmentation entirely.

### Disruption of photoreceptors by early dechorionation

Immunostaining with an anti-Arrestin antibody labels the Group I and II photoreceptor cell bodies and their axons (Horie et al., 2005). The axons are observed as a bundle projecting to the posterior brain vesicle (Figure 2G, white arrow), where they synapse to relay interneurons (Ryan et al., 2016). This pattern of Arrestin staining was observed in all late-dechorionated larvae examined (n=12; Figure 2G and K). By contrast, nearly all early-dechorionated larvae had highly abnormal Arrestin staining patterns (14 of 23), or no recognizable photoreceptors at all (5/19). This included apparent left-right reversals (7 of 23) (Figure 2H and L), or irregular or loosely structured photoreceptor clusters, and with many no longer showing the characteristic close apposition to the ocellus pigment cell (Figs. 2G-J). For example, the larva shown in Figure 2I has two well separated clusters of photoreceptors, each associated with a pigment cell. Moreover, the projections from these photoreceptors project much more posteriorly than that they do in control larvae (red arrow), indicating that the photoreceptor connectivity in the early-dechorionated larvae can be highly irregular. Figure 2L shows an enlargement of a larva with a left-right mirror inversion, but while most of the photoreceptors project correctly (white asterisk), some photoreceptors appear to have missed their pBV targets and instead project too posteriorly (red arrow). We also observed consistent staining of palps (*pp* in Figure 2) with the Arrestin antibody in early-dechorionated larvae, but not in late-dechorionated larvae or non-dechorionated larvae (Horie et al., 2005); the reason for this is not clear.

### Antenna cells and coronet cells

We also assayed for the presence of the paired antenna cells and the coronet cells, as these sensory organs display a stereotyped asymmetry in the larval body (Figure 1). We used an *in situ* strategy to label these organs. By using the hybridization chain reaction (HCR) *in situ* procedure (Molecular Instruments), two fluorescent probes were imaged simultaneously in the same larva, allowing us to assess correlations in symmetry defects between organs. The glutamatergic antenna cells project from the otolith to relay neurons in the posterior brain vesicle, and can be visualized with probes to the vesicular glutamate transport transcript (VGLUT) (Horie et al., 2008; Kourakis et al., 2019). While VGLUT also labels photoreceptors, palp, and epidermal sensory neurons (ESN), the antenna cells can be readily identified by their shape and position in the brain vesicle. We assessed antenna cells after both early and late dechorionation (Figure 3). In late-dechorionated larvae, the VGLUT label allowed us to identify the antenna cell pair in most larvae (10/11); the antenna cells in these larvae showed the expected asymmetry (Figure 3A). In these larvae the photoreceptors were also identifiable in all larvae, and were found in the expected position on the right side of the BV. In early-dechorionated larvae, by contrast, antenna cells were recognized in 58 percent of the larvae observed (11/19), and when present, many showed sidedness defects, positioned on the right instead of the left side (8 of 11) (Figure 3B and D). In those cases where VGLUT labeling indicated the presence of antenna cells, the cells were always observed as a pair, as would be expected, never as single cells or never more than two. The photoreceptors were absent at nearly the same rate as the antenna cells (9/19), but presence or absence of the one did not necessarily indicate presence or absence of the other (data not shown).

**Figure 3.**
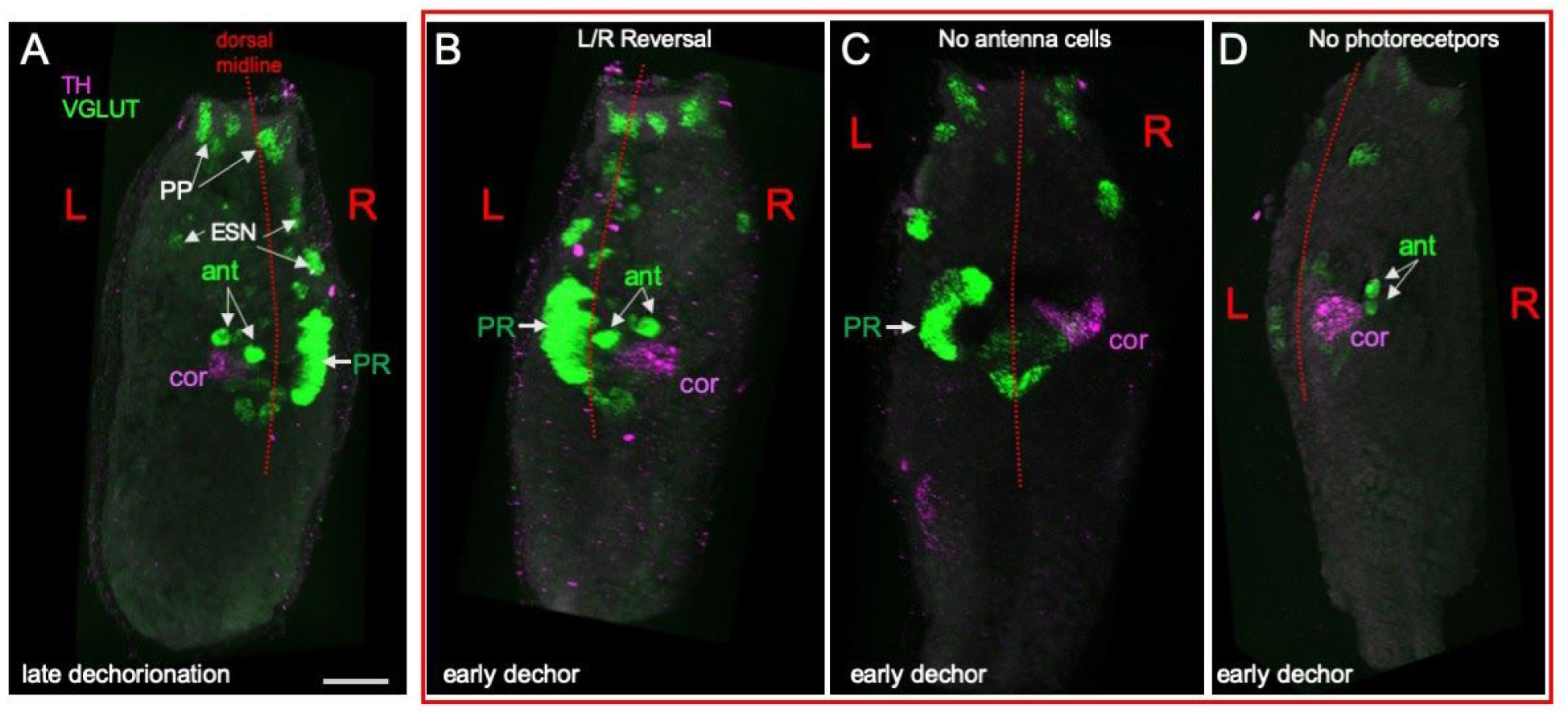
Disruptions in sensory systems caused by early dechorionation. **A**. A late-dechorionated larva showing the paired VGLUT-expressing antenna cells. Also labeled by *in situ* hybridization are the VGLUT-expressing photoreceptors and epidermal sensory neurons, and the tyrosine hydroxylase (TH)-expressing coronet cells. **B-D**. Larvae represent a range of disruptions to sensory systems caused by early dechorionation. Abbreviations: pp:palps; ESN; epidermal sensory neurons; PR: photoreceptors; ant: antenna cell; cor: coronet cell.

The coronet cells of *Ciona* are likely dopaminergic as they label with an *in situ* riboprobe for tyrosine hydroxylase (TH) as well as by anti-dopamine immunolabeling (Moret et al., 2005). A patch of TH-positive cells of the expected size and position (posterior to the antenna cells, and at the left ventral side of the anterior BV) was observed in all late-dechorionated larvae (11/11), as has been described previously (Moret et al., 2005) (Figure 3A). Most early-dechorionated larvae (17/19) also had a TH positive patch in the anterior brain vesicle, indicating the presence of coronet cells. Although the presence of coronet cells seemed robust compared to the antenna cells (and photoreceptor cells), sidedness appeared nearly random, with 7 located to the left of the midline and 10 appearing to the right.

While correlations between presence/absence of antenna cells, coronet cells, and ocellus-associated photoreceptor cells were not apparent in early-dechorionated larvae, sidedness relations did show consistency. The coronet and antenna cells (when both present) were found on the same side. The photoreceptor neuron clusters were found opposite the antenna and coronet cells. That said, in early-dechorionated larvae, VGLUT labeling did not always reveal a coherent group of photoreceptor neurons, as would be seen in typical non-dechorionated larvae. This observation for VGLUT is consistent with the results for the Arrestin antibody (Figure 2) which showed patchy or small clusters of labeling in some early dechorionated larvae.

### Left-right asymmetry in the pBV

The photoreceptors, antenna cells, coronet cells, and a subset of the ESNs all project to the pBV (Figure 1), where they synapse to several classes of interneurons, including the relay neurons, which in turn project posteriorly to the motor ganglion. *In situ* hybridization studies have shown that the pBV made up of two distinct and nested domains, an anterior region of cholinergic neurons (VACHT expressing), and a posterior region of GABAergic neurons (VGAT expressing) (Figure 4A; Kourakis et al., 2019). When viewed laterally, where left-right asymmetry would not be evident, the overall complement and nested anterior-posterior domains of cholinergic and GABAergic neurons appear to be preserved in early-dechorionated larvae (Figure 4B). However, as is evident in the early-dechorionated larva shown in Figure 4B, the angle of the cholinergic and GABAergic neuron clusters with respect to the body axis often differed from the late-dechorionated larvae, suggesting an left-right asymmetry in the pBV.

**Figure 4.**
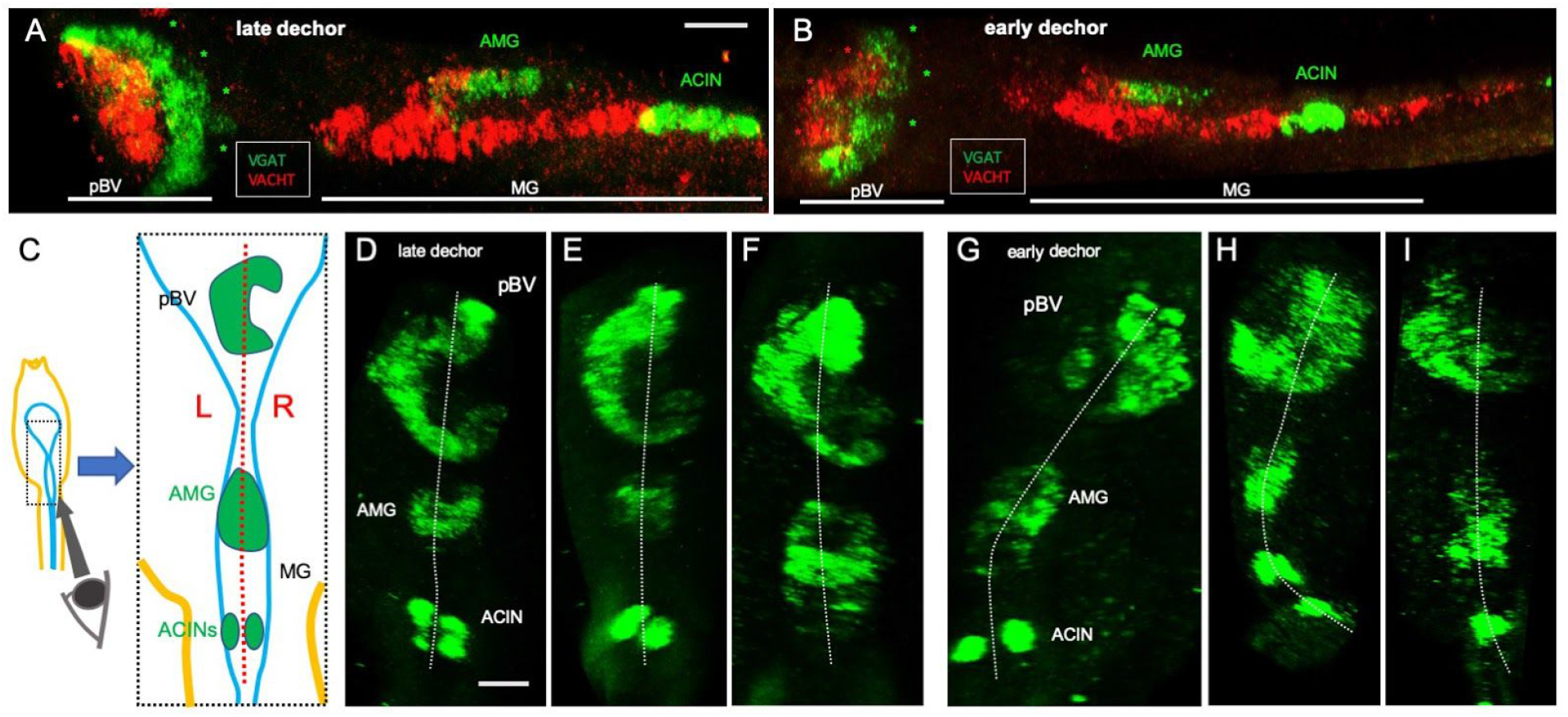
**A** and **B**. Double label *in situ* hybridization for VGAT and VACHT show the similar patterns for both early and late-dechorionated larvae in the layered domains of the pBV (left-lateral view; anterior is left). Also labeled are the VGAT-positive AMG neurons, just dorsal to the VACHT-positive (red) neurons of the MG, and anteriormost set of paired ACINs. **C**. Diagram shows perspective for VGAT *in situs* shown in panels D-I. **D-F**. Late-dechorionated larvae show characteristic right-facing crescent of VGAT expressing cells in the pBV. **G-I**. The asymmetry of the pBV, as seen by VGAT expression, is disrupted in early-dechorionated larvae. Abbreviations: pBV: posterior brain vesicle; MG: motor ganglion; AMG: ascending motor ganglion neurons; ACIN:ascending contralateral inhibitory neurons.

The pBV left-right asymmetry is evident when the VGAT domain is viewed in a transverse plane from the caudal end of the larva (Figure 4C). When viewed from this perspective, the pBV VGAT neurons in late-dechorionated larvae cluster to form a right-facing crescent (Figure 4D-F). In early-dechorionated larvae, the distribution of VGAT neurons in the pBV no longer shows this pattern (Figure 4G-I). Most (4/5) could be interpreted as having a reversed orientation for the VGAT, with the crescent open toward the left. Thus, like the sensory structures in the aBV, the arrangement of interneurons of the pBV has an inherent left-right pattern, and this pattern is disrupted by early dechorionation.

### Asymmetric Pitx in the motor ganglion

Unlike in the brain vesicle, the distribution of neurons in the motor ganglion shows bilateral symmetry. This is observed, for example, in the two cholinergic descending decussating neurons (ddNs) in the anterior MG, which are found paired across the midline, one right and one left. Posterior to the ddNs are right and left pairs of six cholinergic MNs and ten cholinergic MGINs (Figure 1; (Ryan et al., 2016). While some variation has been observed between larvae in the number of glycinergic ascending contralateral inhibitory neurons (ACINs) and MNs, as well as how closely left-right neuron pairs are juxtaposed across the midline in the posterior MG, these asymmetries are rare and appear to occur at random in untreated larvae and thus are likely not driven by the Nodal left-right asymmetry pathway (Kourakis et al., 2019; Ryan et al., 2016).

*Pitx* gene expression, as a downstream target in the Nodal signaling pathway, is diagnostic for left-right asymmetry. In ascidian neurula embryos (*Ciona* and *Halocynthia roretzi*) it is expressed in the left epidermis and includes the the prospective oral siphon primordium by tailbud stages (Boorman and Shimeld, 2002; Christiaen et al., 2002; Morokuma et al., 2002). In late-dechorionated *Ciona* larvae, we observed *Pitx* expression in the left motor ganglion, as well as faintly in the photoreceptors, and in the oral siphon primordium -- a symmetric midline structure (Figure 5A). Expression in the motor ganglion was restricted to its anterior portion, and did not appear to extend to the VGAT-positive (glycinergic) ACIN cells (Figure 5A). The left-right demarcation, however, is very clear, with little or no detectable *Pitx* expression in the right MG. This expression suggests that later *Pitx* expression, like that of neurula and tailbud stages, plays a role in left-right patterning. Moreover, just as early dechorionation disrupts embryonic *Pitx* expression, often resulting in bilateral epidermal expression, early dechorionation likewise disrupted the asymmetric expression of *Pitx* in the MG at the larval stage (Figure 5B). Most early-dechorionated larvae showed bilateral expression of *Pitx* in the MG (4/6 bilateral; 1/6 no expression; 1/6 left-sided); all late-dechorionated controls showed the left-sided pattern (8/8).

**Figure 5.**
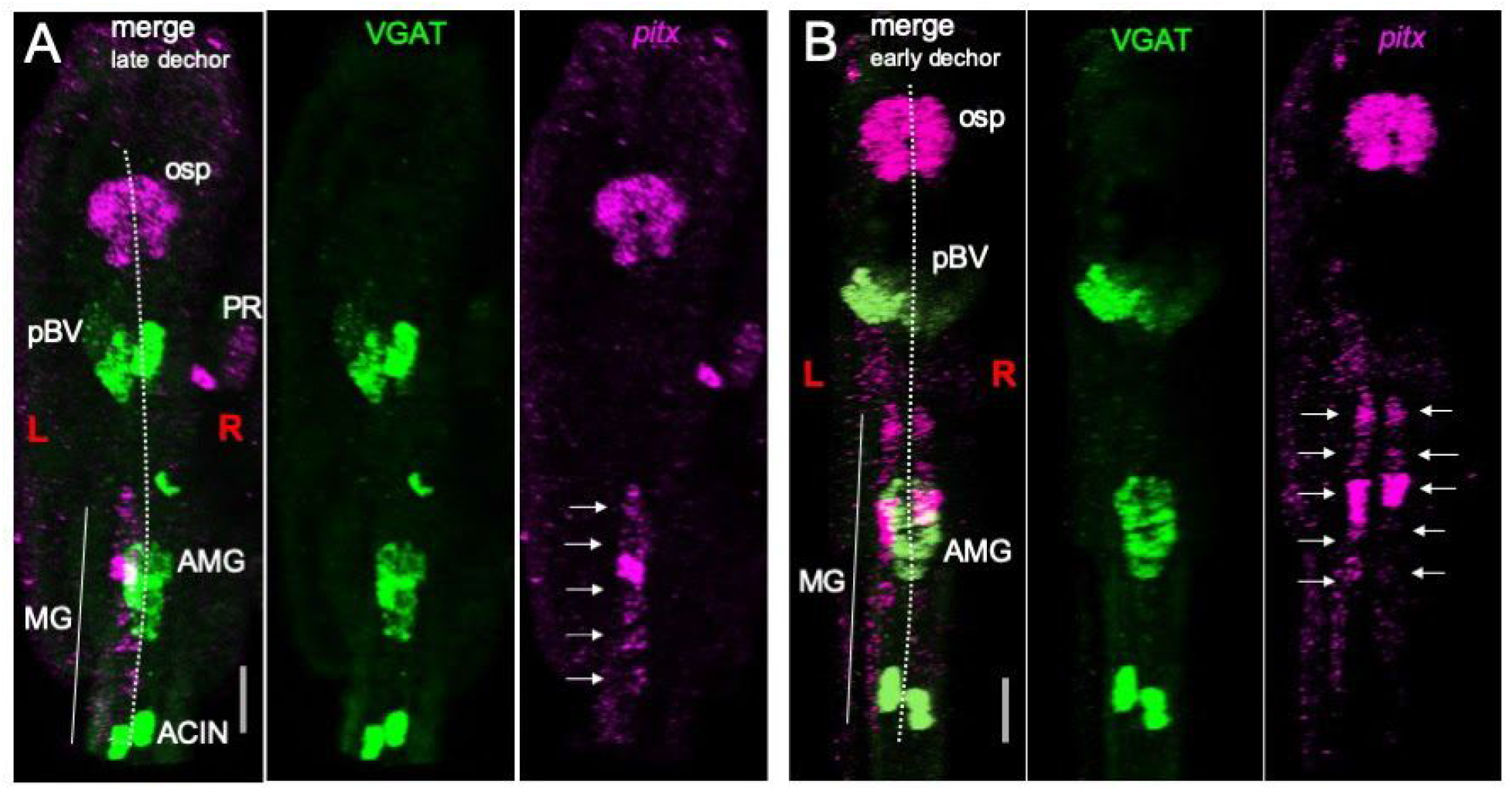
The *Pitx* gene is expressed asymmetrically in larval *Ciona robusta*. **A**. Detail of a late-dechorionated 25 hpf *Ciona* larvae showing *Pitx* expression by *in situ* hybridization co-labeled with VGAT (anterior is up; midline indicated by dotted line). *Pitx* shows left-sided expression in the MG (arrows), in contrast to VGAT which is expressed in the MG on both sides of the midline. *Pitx* expression in the osp is also bilateral. In this view, *Pitx* is also seen in the photoreceptor cells on the right side. **B**. An early-dechorionated larva shows the same bilaterality for VGAT in the MG, and for Pitx in the osp, but *Pitx* in the MG is found on both left and right sides of the dorsal midline (Scale bars are 20 µm). Abbreviations: pBV: posterior brain vesicle; PR: photoreceptor; MG: motor ganglion; AMG: ascending motor ganglion neurons; ACIN:ascending contralateral inhibitory neurons.

A previous study using a transgenic *Pitx-*promoter expression construct and *in situ* probes for Pitx isoforms also reported asymmetric expression in the left motor ganglion at late tail bud and hatching stages (Christiaen et al., 2005). However, these labeling methods required early dechorionation, so it was not clear whether the asymmetric MG expression was typical. Our *in situ* patterns confirm these previous results, showing that left-right asymmetries are present in the MG at larval stages, despite the anatomical left-right symmetry in the number and distribution of MG neurons. Additionally, we can add to asymmetric larval Pitx expression a likely dependence on earlier Nodal-dependent events, since early dechorionation disrupts the left-dominant pattern of Pitx in the MG.

### Left-sided AMPA receptor expression in the motor ganglion interneurons

While the MG approximates bilateral symmetry in its complement of cells, we found that like *Pitx*, a receptor for the neurotransmitter glutamate, the AMPA receptor (AMPAR), is asymmetrically expressed in larval MG. In late-dechorionated *Ciona* larvae *in situ* reveals strong AMPAR expression in cells on the left side of the MG (Figure 6A); expression is also evident anteriorly in the brain vesicle, as reported previously (Kourakis et al., 2019). Cells on the right side of the MG show only faint AMPAR expression. This pattern was observed in 10/10 of late-dechorionated larvae examined. Like *Pitx*-labeled larvae of the same stage, larvae which had been dechorionated early did not show the dominant left-sided expression pattern for AMPAR, consistent with other asymmetries observed in the *Ciona* larval CNS. Some larvae showed reversals, with right-sided expression predominant (4/8; Figure 6B), while others showed bilateral expression (3/8; Figure 6c) and one was observed with the left-dominant pattern.

**Figure 6.**
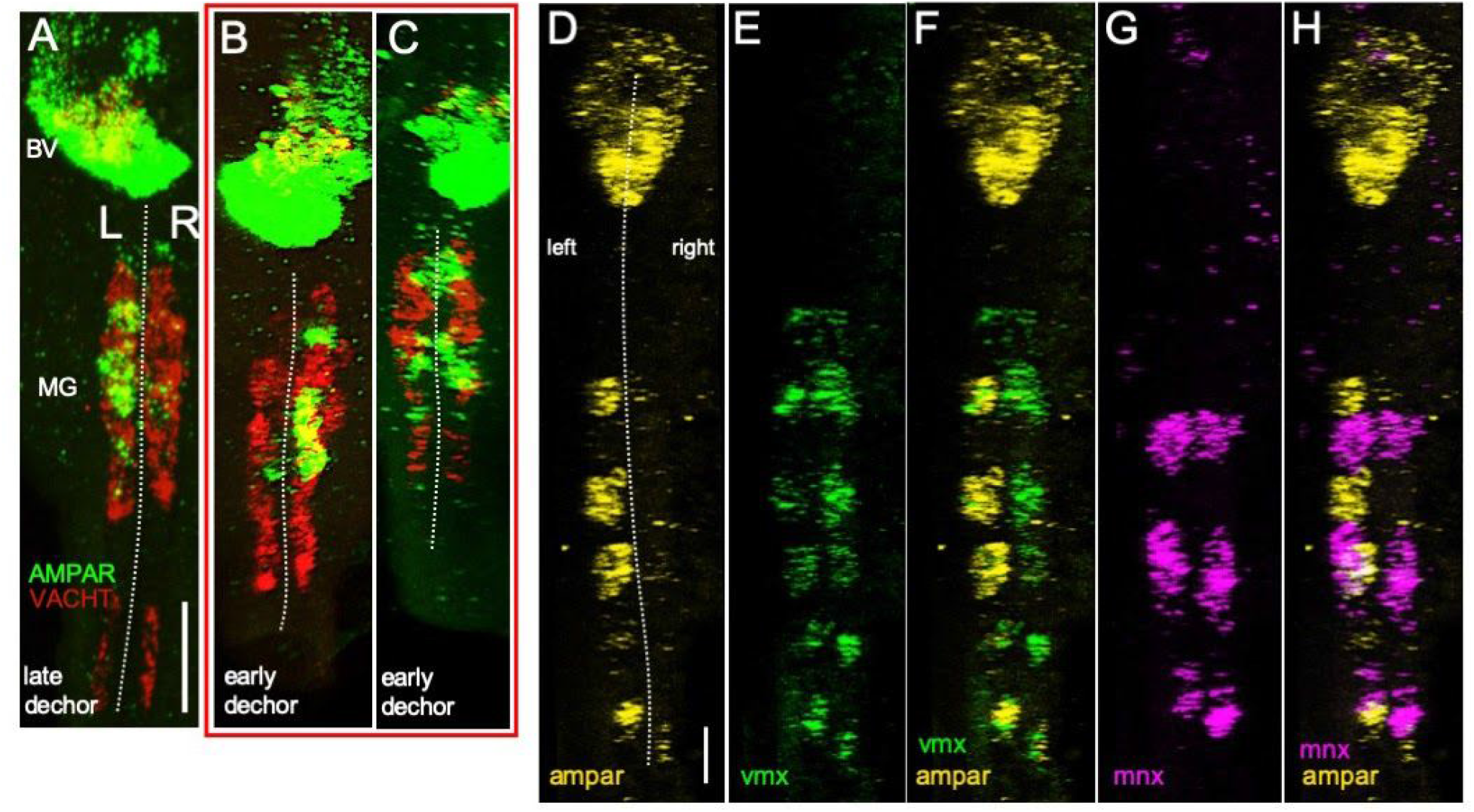
Asymmetric AMPA receptor expression in the Motor Ganglion. **A**. AMPA receptor (AMPAR) is expressed in the left motor ganglion of late-dechorionated larvae. Also shown are VACHT-expressing cells, which are found on both left and right sides of the MG. **B** and **C**. Asymmetric AMPAR expression is disrupted by early dechorionation. Phenotypes include reversals (B) and bilateral expression (C). **D**-**H**. Asymmetric AMPAR is coexpressed with *vmx*, a marker of motor ganglion interneurons, but not *mnx*, a marker of motor neurons. Scale bar in A is 20 µm, and 10 µm in D.

Two of the primary cell classes in the MG are the MNs and the MGINs, which are marked by the expression of *mnx* and *vsx*, respectively (Gibboney et al., 2020). We performed a triple *in situ* with *vsx, mnx* and *AMPAR* to determine which cell classes expressed AMPAR in the MG (Figure 6 D-H). AMPAR shows coexpression with *vsx*, but not *mnx*, indicating it is expressed in the MGINs, and indicted that the MGINs potentially have an asymmetric, left-sided bias in their ability to respond to the neurotransmitter glutamate, suggesting a possible mechanism for asymmetric (*i*.*e*., curved) swim trajectories.

### Disruptions to larval behavior by dechorionation

A strength of the *Ciona* system, which combines a very small and well-documented nervous system with quantifiable behaviors, is the potential for identifying the roles of individual neurons in underlying behavioral control. Unfortunately, many of the techniques that could aid in this analysis, such as transient transgenesis and CRISPR, require early dechorionation. As detailed above, early dechorionation results in profound and variable defects throughout the larval CNS. Consistent with the defects in CNS, we find that the behavior of early-dechorionated larvae, and to a much lesser extent late-dechorionated larvae, is abnormal when compared to non-dechorionated controls.

To assess the behavior of larvae, we used a previously described gravitaxis assay (Bostwick et al., 2020a). In this assay, the behavior of larvae is recorded in a vertically-mounted and covered 6 cm petri dish. In response to a dimming of ambient light, the larvae initiate robust negative gravitaxis swims (i.e., upward swimming). Under constant illumination, the larvae show no up or down bias in their swims. Thus this assay demonstrates the integration of gravity and visual inputs, and gives a measure of the functioning of both the visual and gravitaxis systems. For these assays presented here, which consisted of dimming a 505 nm light from 600 lux to 0 lux (while recording with constant 700 nm illumination), we recorded both the percentage of larvae responding to the dim, which was evident in recordings as swimming behavior initiated immediately following the light dim, as well as the direction of swim trajectories with respect to gravity for those larvae that did respond. All behavior assays were analyzed as described [(Bostwick et al., 2020a), and see Materials and Methods]. The values presented below are the averages and variations (+/- S.D.) between clutches of larvae for each condition. The number (n) represents the total number of larvae analyzed.

While 100% of the non-dechorionated control larvae responded to the dim (n=55), we observed that 83 ± 29% (n=42) of early-dechorionated larvae responded. However, unlike the controls, which displayed a strong bias towards upwards swimming (96.1 ± 3.5% UP; n=55), only 36.2 ± 6.6% of the early-dechorionated larvae (n=27) that responded to the dim swam upwards, while 45.4 ± 7% swam downwards and 18.4 ± 11% swam sideways, illustrating a clear difference between control and early-dechorionated larval behavior (Figure 7A, *p* < 0.00001). Figure 7C (and Movie 1) shows representative swims for control and dechorionated embryos. The swims are indicated by temporal projections of the swims over 20 seconds, which appear as lines. By comparison to early-dechorionated larvae, the behavior of late-dechorionated larvae was more similar to controls. While 97.4 ± 3.6% of our non-dechorionated control larvae (n=140) for this assay responded to the dim, 94.4 ± 7.9% of late-dechorionated larvae (n=61) responded as well. We observed a bias towards upwards swimming in both controls (94.9 ± 0.5% UP; n=136) and late-dechorionated larvae (77.9 ± 8.6% UP; n=57) (Figures 7B and 7C, *p* > 0.05).

**Figure 7.**
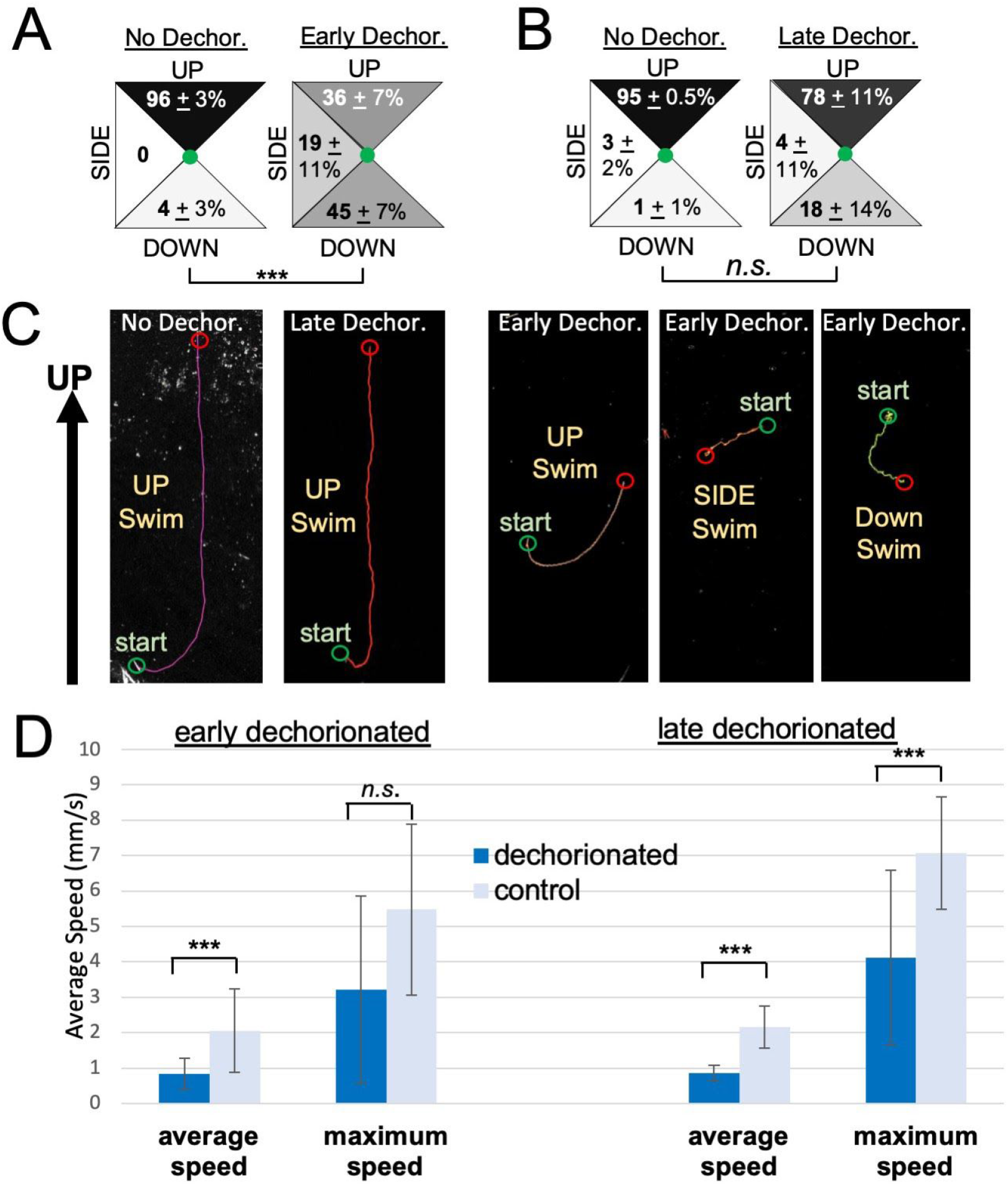
Behavioral disruptions caused by dechorionation. **A**. Early-dechorionated larvae have disrupted negative gravitaxis (upward swimming). **B**. The negative gravitaxis behavior of late-dechorionated larvae more closely resembles non-dechorionated controls than do early-dechorionated larvae. **C**. Representative dimming-induced swims in non-dechorionated, late-dechorionated and early-dechorionated larvae. Lines represent swim trajectories projected across 20 seconds. **D**. Both early and late-dechorionated larvae swim more slowly than do non-dechorionated controls.

In addition, the swimming characteristics of dechorionated larvae were qualitatively different from non-dechorionated control larval swimming (Figure 7D). We observed that the average swimming speed of early-dechorionated larvae (n=13) was 0.84 ± 0.45 mm/s, while the average speed of their controls (n=15) was 2.06 ± 1.18 mm/s (*p*=0.0001). The differences in swim speeds can be observed in Movie 1. Late-dechorionated larvae also tended to swim at slower average speeds than their non-dechorionated controls. The average swimming speed of late-dechorionated larvae (n=11) was observed to be 0.87 ± 0.21 mm/s, while their controls (n=60) swam 2.17 ± 0.59 mm/s on average (*p* < 0.00001). Further, the average maximum speed at which late-dechorionated larvae were able to swim (4.12 ± 2.47 mm/s) was significantly different from the average maximum speed of their non-dechorionated controls (7.06 ± 1.58 mm/s; *p* = 0.0003). This difference in swimming speed is likely not attributable to a different pattern generation for swimming, as no difference was observed between the number of tail beats per second in late-dechorionated larvae (2.35 ± 0.75 flicks/sec; n=12) compared to their controls (2.00 ± 0.53 flicks/sec; n=30; *p* > 0.05). This suggests that the difference in speed might be mechanical, perhaps due to the reduced amount of test in the larvae, making the larvae less rigid. We also did not observe any significant differences between the swimming path tortuosity of early or late-dechorionated larvae and their respective control non-dechorionated larvae (data not shown).

### Selection of early-dechorionated larvae for normal and mirror-image behaviors

In many larvae, disruptions to left-right asymmetry pathways give rise to a range of defects, from mixed or ambiguous asymmetry, to mirror image reversals, and finally to a subset of larvae with apparent normal left-right asymmetry (Levin, 2005). In early-dechorionated *Ciona*, we observed variation in both the number and position of sensory cells (photoreceptors and antennae cells), as well as variation in the shape of cell clusters in the pBV, and in gene expression (*Pitx* and AMPAR) in the MG. However, within all of the early-dechorionated clutches of larvae we assessed, there was usually a small fraction with either apparent normal development, or what appeared to be perfect mirror-image reversals. Likewise, we observed that a small fraction of the early-dechorionated larvae showed the expected upward swimming in response to dimming. Given these observations, we attempted to select early-dechorionated larvae based on their behavior, in order to determine if normal behavior correlated with more normal CNS development. Moreover, larvae selected this way might prove to be useful for assays or protocols requiring early dechorionation.

For the selection procedure, 23-24 hpf larvae were placed in 10 cm agarose-coated petri dishes and observed under a stereomicroscope. The light was briefly dimmed and larvae that responded to the dimming with vigorous upward swimming were picked with a BSA-coated micropipette tip and transferred to a new petri dish (see details in Material and Methods). From approximately 600 dechorionated larvae, we would recover 4-7 larvae with this method in about 15 minutes. At 25 hpf, the behavior of the selected larvae was then reassessed and recorded in the gravitaxis assay, as described above. We observed that 77.2 ± 12% of the selected early-dechorionated larvae (n=23) swam up in response to dimming. By contrast, only 34.3 ± 8.1% of unselected larvae (n=12) swam upwards (Figure 8A). Finally, 96.7 ± 4.7% non-dechorionated control larvae (n=35) upwards after dimming. Thus, we are able to select for larvae whose behavior more closely matches that of the non-dechorionated controls.

**Figure 8.**
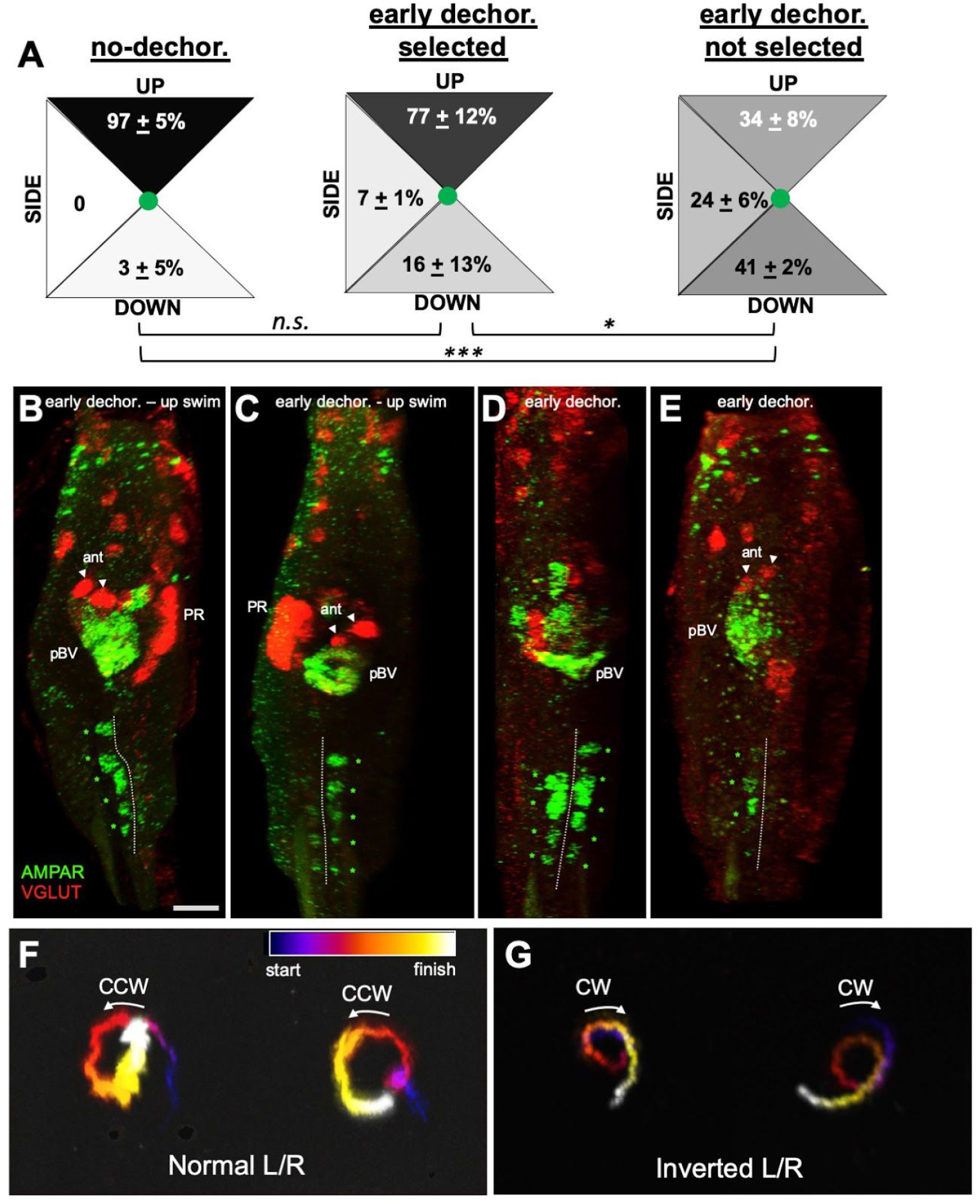
**A**. Gravitaxis of larvae pre-selected for their upward swimming behavior (*selected*) and their non-dechorionated, and not selected siblings. **B-E**. *In situ* hybridization for VGLUTAMPAR of early-dechorionated larvae, either selected for their upward swimming behavior (B and C), or not selected (D and E). **F** and **G**. Temporal projection of swimming trajectories for selected larvae with normal (F) and mirror-image left-right asymmetry (G) following light dimming (projections over 13 seconds). Abbreviations: *: p<0.05; ***: p<0.001; *n*.*s*.: not significant. pBV: posterior brain vesicle. ant.: antenna cells. PR: photoreceptors. L/R: left-right.

The left-right asymmetry of the selected larvae was examined by *in situ* hybridization, using the VGLUT and AMPAR probes, as well as by examining the pigmented cells of the otolith and ocellus. For larvae that had been selected by the behavioral assay, 14 of 33 examined had correctly positioned pigment cells, while 16 had what appeared to be perfect reversals, two had the pigmented cells at the midline, and a single larva was observed with three pigment cells, likely two ocelli and a single otolith pigment cells. By contrast, in non-selected larvae, pigment cell reversals (10 of 20) were more prevalent than the normal pattern (4 of 20) or than midline placement of both otolith and ocellus pigments (6 of 20); a single animal in the latter category showed a split/double ocellus, positioned posteriorly to a single otolith. To further compare selected and non-selected larvae, we used VGLUT *in situ* hybridization to assess two easily identifiable features based on their position and morphology, the asymmetrically placed left-ventral antenna cell pair and right-sided photoreceptors. In non-selected larvae, only three of twenty-three larvae examined showed the expected left-side antenna cells *together with* right-side photoreceptors (13%); one showed a mirror-image reversal with a left-sided photoreceptors and ventral-right antenna cells. Of the remaining nineteen larvae, seven had photoreceptors (whether right or left) without antenna cells (30%), while eleven had identifiable antenna cells, but no photoreceptors (48%). Of thirty-three selected larvae examined, thirty had both photoreceptors and antenna cells (91%), that number evenly split between larvae with normal asymmetry (antenna cells on left, photoreceptors on right) and inverted, mirror-image asymmetry (i.e., a reversal). The remaining three included a single larva with ectopic photoreceptors (this larva also had three pigment cells), one with no recognizable photoreceptors, and one with neither photoreceptors nor antenna cells (though other VGLUT-positive cells were present). A subset of larvae were also cohybridized with the AMPAR probe, which showed that mirror-image reversals in selected larvae determined by VGLUT expression were accompanied by a reversal of AMPAR in the MG (4/4). Larvae with normal asymmetry, by contrast, showed left-sided AMPAR in the MG (5/6), with a single exception showing bilateral AMPAR. These results provided further support that mirror-image reversals are consistent across the CNS.

Because the selected early-dechorionated larvae appeared to consist of an equal mixture of normal and mirror-image left-right reversals, we undertook to determine if the behaviors of these two phenotypes were qualitatively different. *Ciona* larvae show a highly reproducible asymmetric behavior. When viewed from above, rather than from the side as in the gravitaxis assay, the larvae can be seen responding to light dimming with curved or spiraling swims, and the majority of these (>80%) are counterclockwise (Salas et al., 2018). For the assay here, the selected larvae were placed in two groups, normal or inverted, based on the left-right distribution of the pigment cells. We recorded the behavior of six normal and five inverted larvae. All larvae were recorded in three to five dimming assays with 5 minute recovery intervals between each (see Material and Methods), and in total, 19 swims from inverted larvae and 16 swims from normal larvae were recorded. For the larvae with normal left-right pigmentation, 14 of the16 swims were counterclockwise (87%), one was clockwise, and one was unscorable. For the larvae with reversed left-right pigmentation 15 of 19 swims were clockwise (79%), one was counterclockwise, one was straight, and two were unscorable. Representative swims for the two groups are shown in Figure 8F and G. The panels show temporal projections of swims in the 13 seconds following dimming. The projections are presented as heat maps to indicate the starts and finishes of the swims. Representative swims for the two groups are also presented in Movie 3. These results show that larval asymmetry, whether normal or mirror-image reversed, correlates to asymmetric swim behavior, clockwise or counterclockwise, and therefore links this behavior to the patterning which establishes left-right asymmetry earlier in the developing neurula.

## DISCUSSION

The left/right asymmetry of the *Ciona* larval CNS has been well documented (Oonuma et al., 2016; Ryan et al., 2016). The *Ciona* connectome further highlights the left/right asymmetry of the larval CNS (Ryan et al., 2016). This asymmetry is evident both in the localization of specific neuron classes, as well as in their synaptic connectivity. The additional left/right asymmetries described here for *Ciona* larval CNS anatomy, gene expression, synaptic connectivity and behavior all are disrupted by early dechorionation, suggesting all are driven by the common Nodal-dependent symmetry-breaking mechanism. Moreover, the results presented here detail the extent and highly variable nature of the disruptions following early dechorionation.

Given the profound disruptions to CNS anatomy caused by early dechorionation, it is hardly surprising that the behavior of dechorionated larvae is abnormal. What was more unexpected was that within a clutch of dechorionated larvae, individuals with apparently normal CNS anatomy and behavior could be found. Equally unexpected is that larvae with mirror-image reversals could be isolated, and that the behavior of these larvae in a dimming assay also represented a mirror-image reversal with respect to normal larvae. Asymmetric behaviors have been described in a range of metazoans (Güntürkün et al., 2020). Among the chordates, the consequences of disruptions to CNS left-right asymmetry on behavior has been detailed most extensively in zebrafish (Miletto Petrazzini et al., 2020). Notably, zebrafish larvae show a similar response to *Ciona* larvae in response to dimming light -- circular swimming. As a population, zebrafish larvae showed no bias for clockwise or counterclockwise swimming in response to dimming, yet individual larvae in repeated trials show a strong bias towards one direction (Horstick et al., 2020). This acquired asymmetry appears to be related to the Nodal-independent hand preference in humans. However, other asymmetric behaviors in zebrafish are dependent on the Nodal pathway (Barth et al., 2005).

While at the cellular level the MG appears to have left/right symmetry, the expression of *Pitx* and AMPAR is asymmetric. Although asymmetric *Pitx* expression in the motor ganglion has been reported previously (Christiaen et al., 2005), we report here that this asymmetric expression is disrupted by dechorionation. This is also true for the left-sided MG AMPAR expression reported here. The first signs of left-right asymmetry in ascidians is *Pitx* and *nodal* expression in the left epidermis (Boorman and Shimeld, 2002; Morokuma et al., 2002; Shimeld and Levin, 2006). The left epidermal expression in *Ciona* extends into the tailbud stages, and by late tail bud nodal expression is observed in the left brain vesicle and endoderm (Yoshida and Saiga, 2011). It is possible this late tailbud expression could also include the MG precursor, as *pitx* is expressed there (Christiaen et al., 2005). The identity of neurons targeting the AMPAR-expressing MGINs in the MG is unclear. There appear to be no glutamatergic neurons that synapse directly to the MGINs. All VGLUT-positive neurons (*i*.*e*., glutamatergic) in the *Ciona* larva are sensory cells (photoreceptors, antenna cells and epidermal sensory neurons) (Kourakis et al., 2019). None of these synapse on the MGINs (Ryan et al., 2016). Instead, the MGINs receive extensive synaptic input from the GABAergic and cholinergic pBV relay neurons, which themselves receive direct input from the photoreceptor, antenna cell, and coronet sensory cells. However, there are glutamatergic neurons that do project to the MG. These are a subgroup of the ESNs known as the *posterior apical trunk neurons* (pATENs). The pATEN projections enter the MG dorsal to the MGINs where they synapse to AMG neurons. However, the pATENs also appear to make synaptic contact to the basal lamina dorsal to the MG near the AMGs (Ryan et al., 2018). It is possible that the release of glutamate to the basement membrane may be received by the MGINs, as the AMGs and the MGINs directly abut one another. The asymmetric localization of the AMPAR may be responsible for translating glutamate release into asymmetric muscle activation. In fact, stimulation of the ESNs causes counterclockwise circular swimming (Movie 3), which would be predicted from preferential activation of the left flank of muscles. This may represent a pathway for evoking circular swims that is an alternative to, for example, the asymmetric projection of inhibitory antenna cell relay neurons to the MGINs (Bostwick et al., 2020a; Ryan et al., 2016).

Despite the widespread use of methods such as transient transfection by electroporation that require early dechorionation in *Ciona*, and other ascidian species, results from such experiments should be interpreted cautiously, particularly given the highly variable nature of the asymmetry defects observed. Where appropriate, such as for *in situ* hybridization to tailbud or older embryos or larvae, the late dechorionation method described here provides a workable solution to this problem. In addition, the behavior-based selection procedure presented here is a solution when early dechorionation is unavoidable, although it is time-consuming and yields relatively few larvae. We found that the selected larvae appear to be reasonable approximations of normal larvae, as assessed by gene expression, pigmentation and behavior, although half represent mirror-image reversals. We anticipate this procedure will be particularly useful for assessing behavior and CNS function, such as with genetically encoded calcium indicators, in transfected larvae.

## ACKNOWLEDGEMENTS

This work was supported by awards from the NIH (HD038701 and NS103774). We acknowledge the use of the NRI-MCDB Microscopy Facility at UC, Santa Barbara. We thank Kerrianne Ryan for her helpful discussions.

