## Supplementary material for "Left/right asymmetry disruptions and mirror-image reversals to behavior and brain anatomy in *Ciona*": Movie Legends

**Supplemental Movies**

**Movie 1**. Gravitaxis assay for control (no dechorionation), late dechorionated and early dechorionated 25 hour post-fertilization larvae. Movie records 26 seconds of behavior and plays at real time. The light is dimmed at 5.7 seconds (DIM). Circles indicate larvae that exemplify the behavior of the group. Up is to the top of the frame for all movies.

**Movie 2.** Dimming response in isolated larvae with normal and reverse left/right asymmetry. The behavior of two larvae are shown for each type. Movie records 26 seconds of behavior and plays at real time. Dimming occurs at 13 seconds (DIM).

**Movie 3.** Touch response of larva. The larva is touched with a pipet tip in the area of the posterior apical trunk neurons (pATENs). The movie shows 32 second of behavior and plays in real time.
